# Robustness of the neuro-muscular octopaminergic system in the face of cold stress

**DOI:** 10.1101/2022.07.25.501383

**Authors:** Sinan Kaya-Zeeb, Saskia Delac, Lena Wolf, Ana Luiza Marante, Oliver Scherf-Clavel, Markus Thamm

## Abstract

In recent decades, our planet has undergone dramatic environmental changes resulting in the loss of numerous species. This contrasts with species that can adapt quickly to rapidly changing ambient conditions, which require physiological plasticity and must occur rapidly. The Western honeybee (*Apis mellifera*) apparently meets this challenge with remarkable success, as this species is adapted to numerous climates, resulting in an almost worldwide distribution. Here, coordinated individual thermoregulatory activities ensure survival at the colony level and thus the transmission of genetic material. Recently, we showed that shivering thermogenesis, which is critical for honeybee thermoregulation, depends on octopamine signaling. In this study, we tested the hypothesis that the thoracic neuro-muscular octopaminergic system strives for a steady-state equilibrium under cold stress to maintain endogenous thermogenesis. We can show that this applies for both, octopamine provision by flight muscle innervating neurons and octopamine receptor expression in the flight muscles. Additionally, we discovered alternative splicing for *AmOAR*β*2* and our results suggest that at least one isoform is needed to survive cold stress conditions. We assume that the thoracic neuro-muscular octopaminergic system is finely tuned in order to contribute decisively to survival in a changing environment.

## INTRODUCTION

Independence from ambient temperature provides a decisive competitive benefit. Insects, however, are not known to have individual physiological thermostasis as mammals or birds do (Nord and Giroud, 2020). Their activity level is highly dependent on ambient temperatures and although some insects use various strategies (passive or active) to vary their body temperature, they cannot maintain a constant core body temperature (Bartholomew, 1981; Josephson, 1981; Block, 1994; Colinet et al., 2018). Nevertheless, eusociality enables some insect species to achieve thermal homeostasis as a super-organism on the colony level (Kadochová and Frouz, 2013; Stabentheiner et al., 2021). As an eusocial insect, the Western honeybee (*Apis mellifera*) has exactly these characteristics. Several strategies are used to keep the colony at a constant temperature during the breeding season and in winter, which also contributes to the almost worldwide distribution of this species (Simpson, 1961; Seeley and Visscher, 1985; Bujok et al., 2002; Stabentheiner et al., 2003; Wallberg et al., 2014; Buckley et al., 2015; Perez and Aron, 2020; Stabentheiner et al., 2021). For the maintenance of the colony’s thermostasis, the workerbee flight muscles are of exceptional importance. For cooling purposes, foragers collect water which is subsequently evaporated from the combs and thus cooling the hive (Kühnholz and Seeley, 1997). In addition, workerbees fan hot air out of the colony with their wings, which in turn creates an airflow that lets cool air in (Egley and Breed, 2013; Fahrenholz et al., 1989). Heat generation occurs exclusively through muscle tremors of the indirect flight musculature and can therefore be referred to as shivering thermogenesis (Stock, 1999; Stabentheiner et al., 2010). Recently we could show, that workerbee thermogenesis relies on octopamine signaling. The dorsoventral wing elevators (DV) and dorsal-longitudinal wing depressors (DL), which constitute the indirect flight muscles, are innervated by octopaminergic neurons from the mesometathoracic ganglion (MMTG, Kaya-Zeeb et al., 2022). The release of octopamine for the purpose of thermogenesis activates β octopamine receptors, most likely AmOARβ2 (Kaya-Zeeb et al., 2022). With *AmOARα1*, another octopamine receptor gene displays dominant flight muscle expression, but we did not find evidence for an involvement of the corresponding receptor sub-type in thermogenesis, yet. Disturbances of the flight muscle octopaminergic system lead to an impairment of thermogenesis and consequently, the affected bees suffer from hypothermia (Kaya-Zeeb et al., 2022). At the colony level, this would endanger the survival of the colony during winter and in extreme habitats from desserts to high mountains. Moreover, new problems arise due to man-made climate change. Increasing temperature fluctuations in short time periods are to be expected, which can lead to rapid temperature decrease (Van Asch et al., 2013; Walther et al., 2002; Rosenzweig et al., 2008). The intensification of such periods poses a major challenge to physiological processes in general and to thermoregulation in particular.

In this study, we used honeybees as a model system to investigate how the thoracic neuro-muscular octopaminergic system responds to cold stress. We applied cold stress to workerbees to force them into thermogenesis in order to avoid chill coma. Subsequently, we determined the concentration of octopamine as well as the expression of the relevant octopamine receptor genes. With this approach we tested the hypothesis, that octopamine is always provided in sufficient quantity by the neurons innervating the flight muscle and, when released, can be recognized by specific receptors. This requires that the expression of receptor genes (at least *AmOARβ2*) is maintained at a constant level or increased as needed.

## MATERIALS AND METHODS

### Cold stress exposure

Brood combs and adult worker honeybees (*Apis mellifera carnica*) were collected from departmental bee colonies. For age-controlled bees (one week old), hatching bees were collected from a brood comb, color-marked and reinserted into a standard hive. Alternatively, newly emerged bees were held in cages (T = 34 °C, RH = 65 %) and fed *ad libitum* with sucrose solution (30 % w/v). After one week, both groups were collected from the hive or the incubator for further experiments. As forager bees, we defined bees that returned to the hive with pollen loads on their hind legs. Prior to each cold stress experiment, bees were subjected a one (1 week old bees) or a two day (forager bees) run-in period with identical conditions (T= 34 °C, RH = 65 %). During the run-in period and the stress conditions, all groups were provided with sucrose solution (30 % w/v) *ad libitum*. All bees that were exposed to stress conditions were immediately flash-frozen in liquid nitrogen after experiencing cold stress conditions and subsequently stored at −80 °C until further experimental procedure (gene expression analysis, monoamine quantification).

### OA injection

Additionally to cold stress, one week old bees (see above) were treated with OA injections. Individual bees were chilled on ice until no further movement could be detected. Then, the thorax was punctured centrally to either inject 1.0 µL saline solution (270 mM sodium chloride, 3.2 mM potassium chloride, 1.2 mM calcium chloride, 10 mM magnesium chloride, 10 mM 3-(N-morpholino) propanesulfonic acid, pH = 7.4; Erber and Kloppenburg 1995) or 1.0 µL octopamine solution (0.01 M in saline, Sigma-Aldrich) using a 10.0 µL Hamilton syringe. The different treatments were kept in different cages and were incubated for either 30 min or 120 min (T = 34 °C, RH = 65 %). Subsequently, all bees were flash-frozen in liquid nitrogen and stored at -80 °C until further procedure.

### Time-survival analysis (Kaplan-Meier estimator)

One week old bees from hive or cage (see above) were transferred to cages of 30 bees each (with sucrose solution (30 % w/v) *ad libitum*). One cage (control) was held under defined conditions (T = 34 °C, RH = 65 %). Two additional cages were exposed to cold stress (10 °C, RH = 65 %). All cages were filmed for 120 min (LifeCam Cinema HD, Microsoft, Redmond, USA) to determine if and when the bees entered chill coma. Afterwards, all the cages in which chill coma occurred were transferred warm conditions to calculate a recovery rate (percentage of bees brought back from the chill coma).

### Monoamine quantification (HPLC-ECD)

For each individual, DV and DL were dissected under liquid nitrogen and merged as one sample. The remaining thoracic tissue was thawed in ice-cold ethanol to undergo immediate dissection of the MMTG. All extracted tissues were kept at −80 °C until extraction of monoamines. The high-performance liquid chromatography (HPLC) protocol represents a slightly modified version as described earlier (Kaya-Zeeb et al., 2022). All samples (tissues and calibrators) were processed and treated equally. We used 120 µL (DV+DL) or 60 µL (MMTG) of the extraction solution (10.0 pg/µL 3,4-dihydroxy-benzylamine (DHBA) in 0.2 M perchloric acid). The raw data was processed with Chromeleon (7.2.10, Thermo Fisher Scientific, Waltham, USA) for further statistical (see below).

### Protein quantification (Bradford)

The pellet from the monoamine extraction was used for protein quantification in oder to normalize monoamine concentrations. After resuspension in 500 µL (DV+DL) or 30 µL (MMTG) NaOH (0.2 M) samples were incubated on ice for 15 min and then centrifuged (9391 g, 5 min). The supernatant (DV+DL: 5 µL, MMTG: 10 µL) were transferred into 1 mL 1x ROTI^®^Nanoquant solution (Carl Roth, Karlsruhe, Germany). All samples and an external calibrator (1, 2, 3, 5, 10, 20 µg/mL Albumin Fraction V; Carl Roth, Karlsruhe, Germany) were analyzed using an Infinite 200 Pro (Tecan, Männedorf, Switzerland).

### Verification of AmOARβ2 isoforms

Sequence analysis was performed with cDNA from thoraces of 20 individual workerbees. Two individual thoraces were pooled in each case and homogenized using 1 ml peqGOLD TriFast™ (Peqlab) following a 5 min incubation at room temperature. After adding of 200 µL chloroform and phase separation, the aqueous phase was transferred to 500 µL isopropyl alcohol. After centrifugation (21,130 g, 15 min, 4 °C), the RNA pellet was purified by two consecutive washing steps (1 ml, 75 % (v/v) ethanol), dried, and diluted in 50 µL RNase-free water. To ensure complete DNA removal we performed DNase treatment (DNase I M0303S; New England Biolabs, Ipswich, Massachusetts) according to the manufacturer’s protocol with subsequent phenol-chloroform extraction (Roti^®^-Aqua-P/C/I; Carl Roth, Karlsruhe, Germany). The purified RNA pellet was diluted in 50 µL RNase-free water and served as template for cDNA synthesis (AccuScript™ High Fidelity 1st Strand cDNA Synthesis; Agilent Technologies, Santa Clara, USA). PCR amplification was conducted using Phusion^®^ DNA Polymerase (New England Biolabs, Ipswich, Massachusetts) on cDNA with *AmOARβ2* isoform (Table 1). After successful amplification and A-tailing (*Taq* DNA Polymerase with ThermoPol^®^ Buffer, New England Biolabs, Ipswich, Massachusetts) the PCR products were subjected to T/A cloning (pGEM^®^-T Vector Systems; Promega, Fitchburg, USA) with subsequent Sanger sequencing (Sanger et al. 1977; Genewiz, Leipzig, Germany).

**Table 1.**
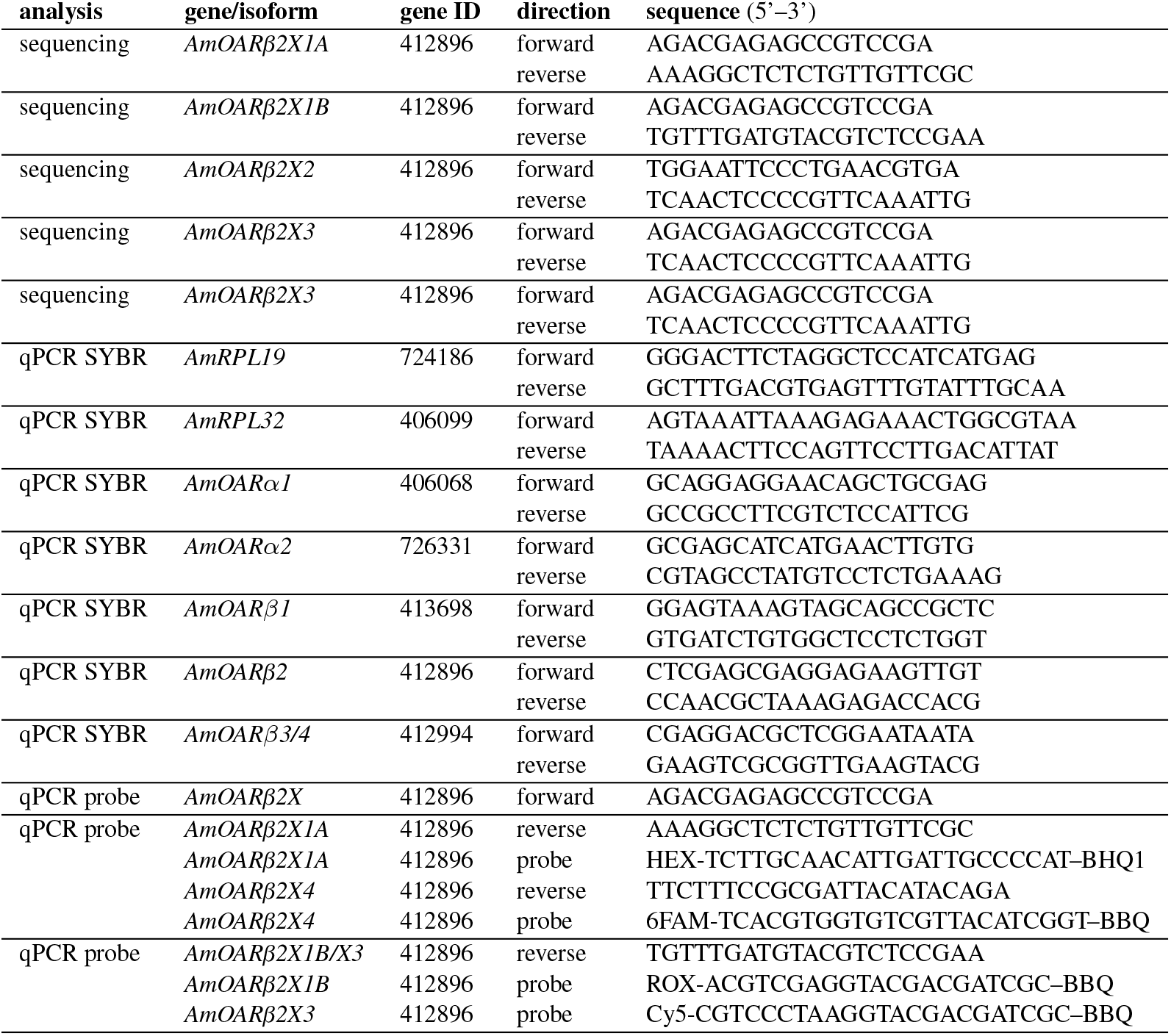
Oligonucleotides used in this study.

### Gene expression analysis

Individual flight muscles (DV+DL) were dissected under liquid nitrogen and stored at -80 °C until further processing. Following the manufacturers standard protocol, we extracted total RNA of individually pooled DV and DL using the GenUP Total RNA Kit (biotechrabbit, Henningsdorf, Germany). We included a DNase I digestion step. After the RNA was bound to the *Mini Filter RNA*, we added 50 µL DNase mix containing 30 U RNase-free DNase I (Lucigen, Middleton, USA) and incubated for 15 min at room temperature. The RNA_total_ concentration was determined photometrically and individual flight muscle cDNA was generated using Biozym cDNA Synthesis Kit (Biozym, Hessisch Oldendorf, Germany). Afterwards, quantitative real-time PCR (qPCR) was used to determine gene expression. Individual cDNAs were analyzed in triplicates per gene. Total reaction volume was 20 µL and contained 4.2 µL H_2_O, 10 µL 2 x qPCR S’Green BlueMix (Biozym, Hessisch Oldendorf, Germany), 0.4 µL of gene specific forward and reverse primer (0.2 µM, Table 1) and 5 µL cDNA. For the analysis of the AmOARβ2 isoforms, we established TaqMan^®^ based duplex assay. Each reaction (20 µL) contained 2.6 µL H_2_O, 10 µL 2 x qPCR S’Green BlueMix (Biozym, Hessisch Oldendorf, Germany), 0.4 µL of each primer (0.2 µM, Table 1), 0.4 µL of each TaqMan^®^ probe (0.1 µM, Table 1) and 5 µL cDNA. All qPCR runs were performed on a Rotor-Gene Q (Qiagen, Hilden, Germany) with following cycling conditions: 95 °C for 2 min, 35 cycles at 95 °C for 5 s and 30 °C for 30 s; followed by a melting curve analysis (not for TaqMan^®^ assays). Gene of interest expression was quantified relative to *AmRPL32* and *AmRPL19* (Kaya-Zeeb et al., 2022; Lourenço et al., 2008).

### Statistical analysis

Statistical analyses were computed using R (4.2.0, R Core Team, 2021) and the R packages ‘car’ (3.0.13, Fox and Weisberg, 2019), ‘dplyr’ (1.0.9, Wickham et al., 2022), ‘FSA’ (0.9.3, Ogle et al., 2022), ‘Rcpp’ (1.0.8.3, Eddelbuettel and François, 2011), ‘reshape2’ (1.4.4, Wickham, 2007), ‘Rmisc’ (1.5.1, Hope, 2022), ‘rstatix’ (0.7.0, Kassambara, 2021), ‘tidyr’ (1.2.0, Wickham and Girlich, 2022), and ‘xtable’ (1.8.4, Dahl et al., 2019). Shapiro-Wilk testing was performed to check for normal distributions. Depending on whether the data were normally distributed or not, they were analyzed using a *t*-test or Mann-Whitney *U* test, respectively. For the analysis of the Kaplan-Meier estimator we additionally used the R packages ‘survival’ (3.3.1, Therneau, 2022; Terry M. Therneau and Patricia M. Grambsch, 2000) and ‘survminer’ (0.4.9, Kassambara et al., 2021). Octopamine receptor gene expressions and *AmOARβ2* isoform abundance were relatively quantified using the R package ‘EasyqpcR’ (1.22.1, Le Pape, 2012). Data was visualized using the R packages ‘cowplot’ (1.1.1, Wilke, 2020), ‘ggplot2’ (3.3.6, Wickham, 2016), ‘ggpol’ (0.0.7,Tiedemann, 2020), ‘ggpubr’ (0.4.0, Kassambara, 2020), ‘grid’ (4.2.0, R Core Team, 2021), ‘ggsignif’ (0.6.3, Constantin and Patil, 2021), ‘magick’ (2.7.3, Ooms, 2021), ‘mdthemes’ (0.1.0, Neitmann, 2020), and ‘png’ (0.1.7, Urbanek, 2013).

## RESULTS

### Responses of workerbees to cold exposure under laboratory conditions

We wanted to analyze the effects of cold stress on workerbees under laboratory conditions, in order to control as many factors as possible. Since our focus was on thermogenesis, these bees additionally should be able to avoid chill coma by active thermogenesis. To our surprise, the one week old bees from cages could not meet these conditions. All bees of this group were found in chill coma latest after 90 min (Figure 1A). The workerbees from the hives (one week old bees, forager bees), on the other hand, withstood the cold without difficulty (Figure 1A). These bees organize themselves into a cluster to effectively perform thermogenesis (Figure 1B). The one week old bees from cages might be in a physiological status that does not enable them to perform thermogenesis. This is supported by the fact that the controls have very low RNA_total_ concentrations in their flight muscles. This is enormously increased by cold stress (Figure 1C). However, this is apparently not sufficient to enable thermogenesis. In hive bees, we see less pronounced but still significant differences in flight muscle RNA_total_ concentration between control and cold-stressed bees (Figure 1C). From these results we conclude the following. First, one week old bees from cages are not suitable for laboratory studies of cold stress and are therefore excluded from further analysis. Second, irrespective whether thermogenesis is actively performed or not, cold stress triggers an increase in transcription activity.

**Figure 1.**
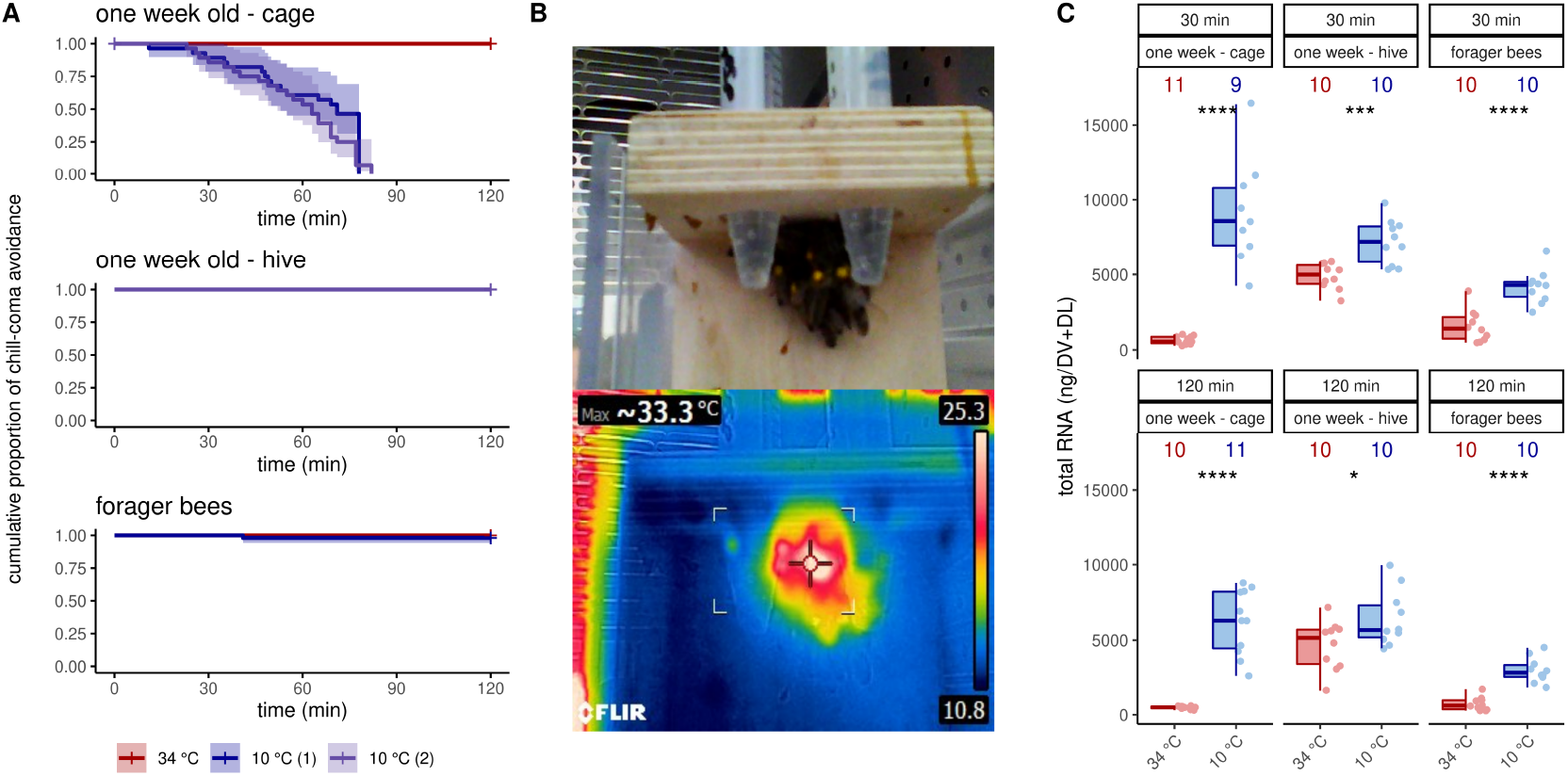
Workerbees respond differently to cold stress. **(A)** One week old workerbees from cages are not able to withstand two hours of cold stress without falling into chill coma (log rank test χ2(2) = 76.1, *p ≤* 0.0001, *N*_34 °C_ = 32, *N*_10 °C (1)_ = 30, *N*_10 °C (2)_ = 31). Not a single one week old workerbees from hives went into chill coma (*N*_34 °C_ = 31, *N*_10 °C (1)_ = 30, *N*_10 °C (2)_ = 30). When forager bees were cold stressed over 120 min, only one bee felt into chill coma (log rank test χ2(1) = 1, *p* = 0.3, *N*_34 °C_ = 48, *N*_10 °C_ = 50). **(B)** Workerbees that successfully survive two hours of cold stress form a cluster in which they effectively perform thermogenesis. **(C)** The RNA_total_ amount significantly increases in workerbee flight muscles due to cold stress (*t*-test, one week - cage - 30 min: *t*(8.07) = -7.20, *p* = 0.0005; one week - cage - 120 min: *t*(10) = -8.84, *p* = 0.035; one week - hive - 30 min: *t*(14.3) = -4.26, *p* = 0.0005; one week - hive - 120 min: *t*(17.7) = -2.28, *p* = 0.035; forager - 30 min: *t*(17.9) = -5.30, *p ≤* 0.0001; forager - 120 min: *t*(14.3) = -7.53, *p ≤* 0.0001). For each group/data-set median ± interquartile range (IQR, left part) and individual data points (right part) are shown.

### Octopamine receptor gene expression

We next asked if this increase in transcription activity also affects octopamine receptor gene expression. In one week old bees, significant differences can only be detected for *AmOARβ2* after 120 min and for *AmOARβ3/4* after 30 min cold stress (Figure 2A, Table 2). For all remaining genes and time points we observed no expression differences (Figure 2A, Table 2). Additionally, we detected no change in the expression of octopamine receptor genes in forager bees after either 30 or 120 min of cold stress (Figure 2B, Table 2).

**Table 2.** Results of the statistical analysis (Mann-Whitney *U* test) of octopamine receptor gene expression in workerbee flight muscles under cold stress (34 °C vs. 10 °C). For visualization please see Figure 2 and Figure 3. * = p *<* 0.05, ** = p *<* 0.01, ns = p *≥* 0.05.

| bees | time | gene/isoform | $N_{34\text{ °C}}$ | $N_{10\text{ °C}}$ | *U* | *p* | *z* | *r* | |
| --- | --- | --- | --- | --- | --- | --- | --- | --- | --- |
| one week old bees | 30 min | *AmOARα1* | 10 | 10 | 47 | 0.853 | -0.185 | -0.041 | ns |
| one week old bees | 30 min | *AmOARβ1* | 10 | 10 | 73 | 0.089 | -1.70 | -0.380 | ns |
| one week old bees | 30 min | *AmOARβ2* | 10 | 10 | 44 | 0.684 | -0.407 | -0.091 | ns |
| one week old bees | 30 min | *AmOARβ3/4* | 10 | 10 | 9 | 0.001 | -3.28 | -0.733 | ** |
| one week old bees | 120 min | *AmOARα1* | 10 | 10 | 49 | 0.971 | -0.036 | -0.008 | ns |
| one week old bees | 120 min | *AmOARβ1* | 10 | 10 | 58 | 0.579 | -0.555 | -0.124 | ns |
| one week old bees | 120 min | *AmOARβ2* | 10 | 10 | 22 | 0.036 | -2.10 | -0.470 | * |
| one week old bees | 120 min | *AmOARβ3/4* | 10 | 10 | 45 | 0.739 | -0.333 | -0.075 | ns |
| forager bees | 30 min | *AmOARα1* | 9 | 10 | 28 | 0.182 | -1.33 | -0.306 | ns |
| forager bees | 30 min | *AmOARβ1* | 9 | 10 | 39 | 0.661 | -0.439 | -0.101 | ns |
| forager bees | 30 min | *AmOARβ2* | 9 | 10 | 30 | 0.243 | -1.17 | -0.268 | ns |
| forager bees | 30 min | *AmOARβ3/4* | 9 | 10 | 44 | 0.968 | -0.04 | -0.009 | ns |
| forager bees | 120 min | *AmOARα1* | 10 | 9 | 39 | 0.661 | -0.439 | -0.101 | ns |
| forager bees | 120 min | *AmOARβ1* | 10 | 9 | 61 | 0.211 | -1.25 | -0.287 | ns |
| forager bees | 120 min | *AmOARβ2* | 10 | 9 | 48 | 0.842 | -0.199 | -0.046 | ns |
| forager bees | 120 min | *AmOARβ3/4* | 10 | 9 | 68 | 0.065 | -1.84 | -0.423 | ns |
| one week old | 30 min | *AmOARβ2X1A* | 10 | 10 | 22 | 0.036 | -2.10 | -0.470 | * |
| one week old | 30 min | *AmOARβ2X1B* | 10 | 10 | 47 | 0.853 | -0.185 | -0.041 | ns |
| one week old bees | 30 min | *AmOARβ2X3* | 10 | 10 | 21 | 0.029 | -2.19 | -0.489 | * |
| one week old bees | 120 min | *AmOARβ2X1A* | 10 | 10 | 17 | 0.012 | -2.53 | -0.565 | * |
| one week old bees | 120 min | *AmOARβ2X1B* | 10 | 10 | 58 | 0.579 | -0.555 | -0.124 | ns |
| one week old bees | 120 min | *AmOARβ2X3* | 10 | 10 | 30 | 0.143 | -1.46 | -0.328 | ns |
| forager bees | 30 min | *AmOARβ2X1A* | 9 | 10 | 42 | 0.842 | -0.199 | -0.046 | ns |
| forager bees | 30 min | *AmOARβ2X1B* | 9 | 10 | 40 | 0.72 | -0.358 | -0.082 | ns |
| forager bees | 30 min | *AmOARβ2X3* | 9 | 10 | 29 | 0.211 | -1.25 | -0.287 | ns |
| forager bees | 120 min | *AmOARβ2X1A* | 10 | 9 | 51 | 0.661 | -0.439 | -0.101 | ns |
| forager bees | 120 min | *AmOARβ2X1B* | 10 | 9 | 76 | 0.010 | -2.57 | -0.59 | * |
| forager bees | 120 min | *AmOARβ2X3* | 10 | 9 | 75 | 0.013 | -2.48 | -0.568 | * |

**Figure 2.**
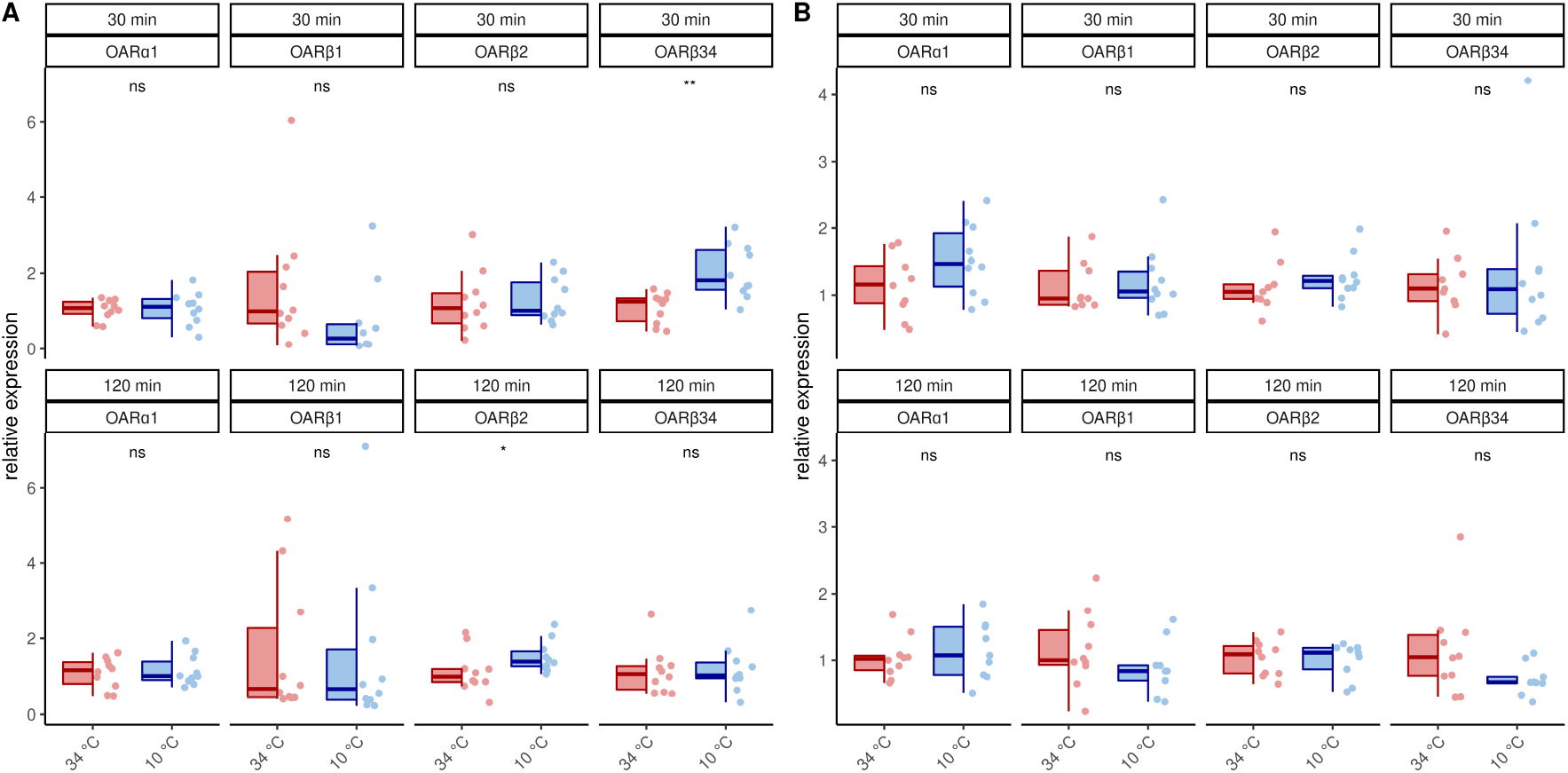
Octopamine receptor gene expression in the flight muscles workerbees under cold stress. One week old workerbees (A) and forager bees (B) were cold stressed for 30 and 120 min and subsequently, the relative octopamine receptor gene expression was quantified in flight muscles (DV+DL). * = p *<* 0.05,** = p *<* 0.01, ns = p *≥* 0.05, Mann-Whitney *U* test. For detailed statistics please see Table 2. For each group/data-set median *±* IQR (left part) and individual data points (right part) are shown.

### AmOARβ2 is spliced alternatively

Publicly available transcriptome data indicate that *AmOARβ2* expresses multiple isoforms due to differential splicing (NCBI, 2022) and we wanted to know, if AmOARβ2 isoform abundance is affected by cold stress. At least five *AmOARβ2* isoforms exist, that differ with respect to their coding sequence (Figure 3A). Both, *AmOARβ2×1A* and *AmOARβ2×1B*, encode an identical receptor protein already functionally described by Balfanz et al. (2013). *AmOARβ2×2* encodes a truncated receptor protein with two trans-membrane domains missing. Likely, this results in impaired integrity as well as poor stability of the receptor protein (Zhu and Wess, 1998; Wise, 2012). We considered it to be unlikely, that a functional receptor protein providing correct ligand interactions could arise from such an isoform and consequently excluded *AmOARβ2×2* from further analysis. Finally we detected *AmOARβ2×3* and *AmOARβ2×4*. Both isoforms encode putative proteins that differ from each other as well as from AmOARβ2×1 by their unique C-terminus sequences.

**Figure 3.**
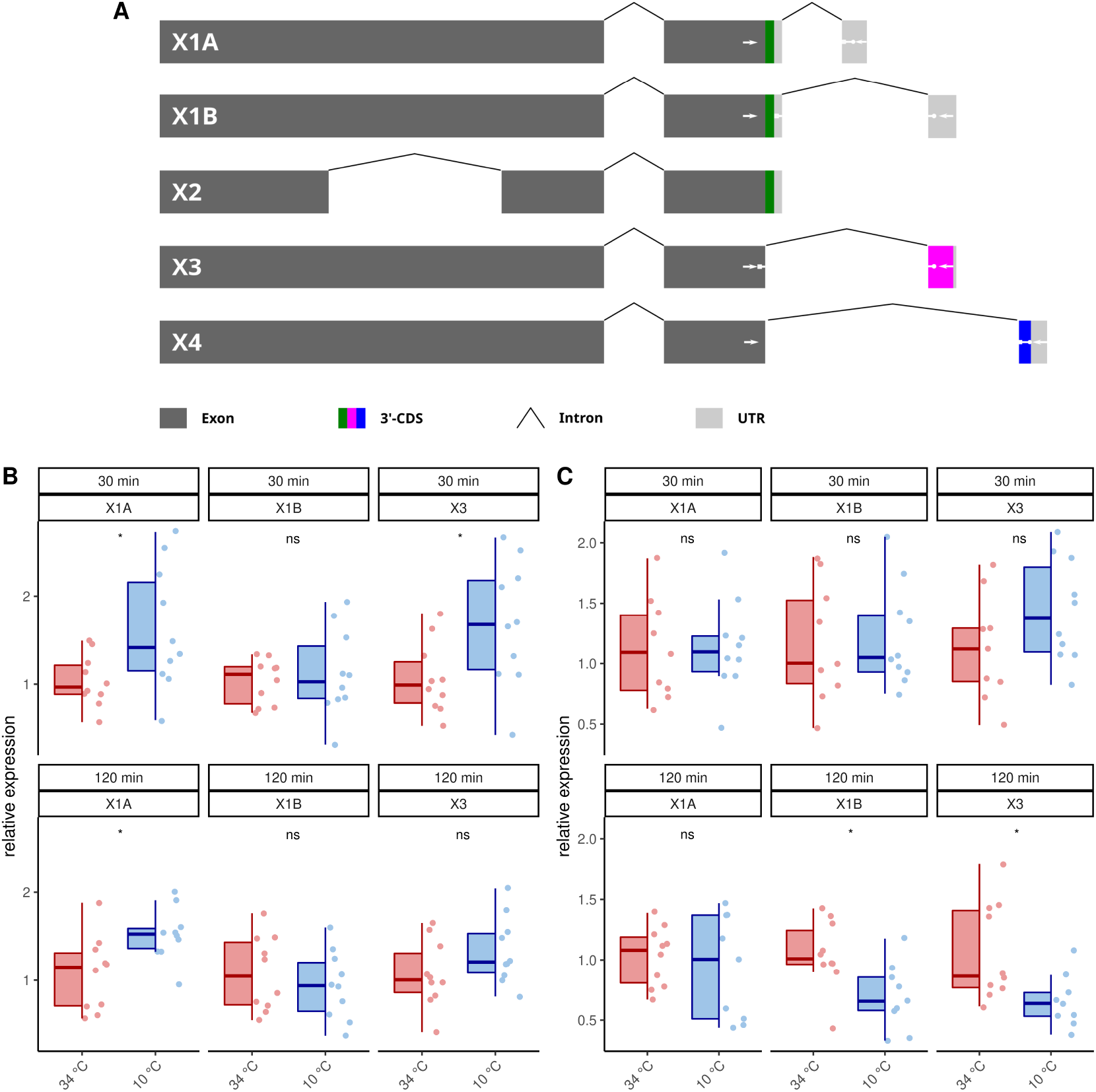
Differential splicing of *AmOARβ2*. (**A**) Schematic representation of *AmOARβ2* alternative splicing. Rectangles in dark gray represent the exons and the introns are visualized by chevron shapes. In multi color the 3’-coding sequence (3’-CDS, including the stop codon) is shown of each isoform. The untranslated region (UTR) is illustrated in light gray. White arrows demark sequences which get specifically targeted by the designed primers and probes (see also Table 1). (**B-C**) Quantification of the *AmOARβ2* isoform abundance in one week old workerbees (B) and forager bees (C). For each group/data-set median*±* IQR (left part) and individual data points (right part) are shown. * = p *<* 0.05, ns = p*≥* 0.05, Mann-Whitney *U* test. For detailed statistics please see Table 2.

Because our standard qPCR assay cannot distinguish different isoforms of this gene, we developed a hybridization probe based assay to analyze the *AmOARβ2* isoform abundance with respect to cold stress (Figure 3B-C, Table 2). *AmOARβ2×1A* abundance is significantly increased after 30 and 12 min cold stress in one week old bees but not in forager bees. No differences can be observed for *AmOARβ2×1B* in one week old workerbees (both time points) and in forager bees after 30 min. After 120 min *AmOARβ2×1B* is significantly decreased in forager bees. *AmOARβ2×3* abundance in increased after 30 min but not after 120 min in one week old bees. In forager bees, no differences can be observed this isoform after 30 min whereas it is reduced after 120 min of cold stress. *AmOARβ2×4* does not reach the threshold for the most individuals and therefore was excluded from analysis.

### Octopamine concentrations are stable under cold stress conditions

In addition to octopamine receptor expression, of course, octopamine is also required for thermogenesis (Kaya-Zeeb et al., 2022). We therefore wondered whether an increased thermogenic activity would require increased octopamine provision. With one exception, cold stress did not change octopamine concentrations in the flight muscles and MMTG (Figure 4). Only forager bees display higher octopamine titers in their flight muscles after 30 min at 10 °C (Figure 4). Besides octopamine, we additionally detected dopamine in the flight muscles and dopamine and serotonin in the MMTG. No differences were found here either (Table S1, Figure S1).

**Figure 4.**
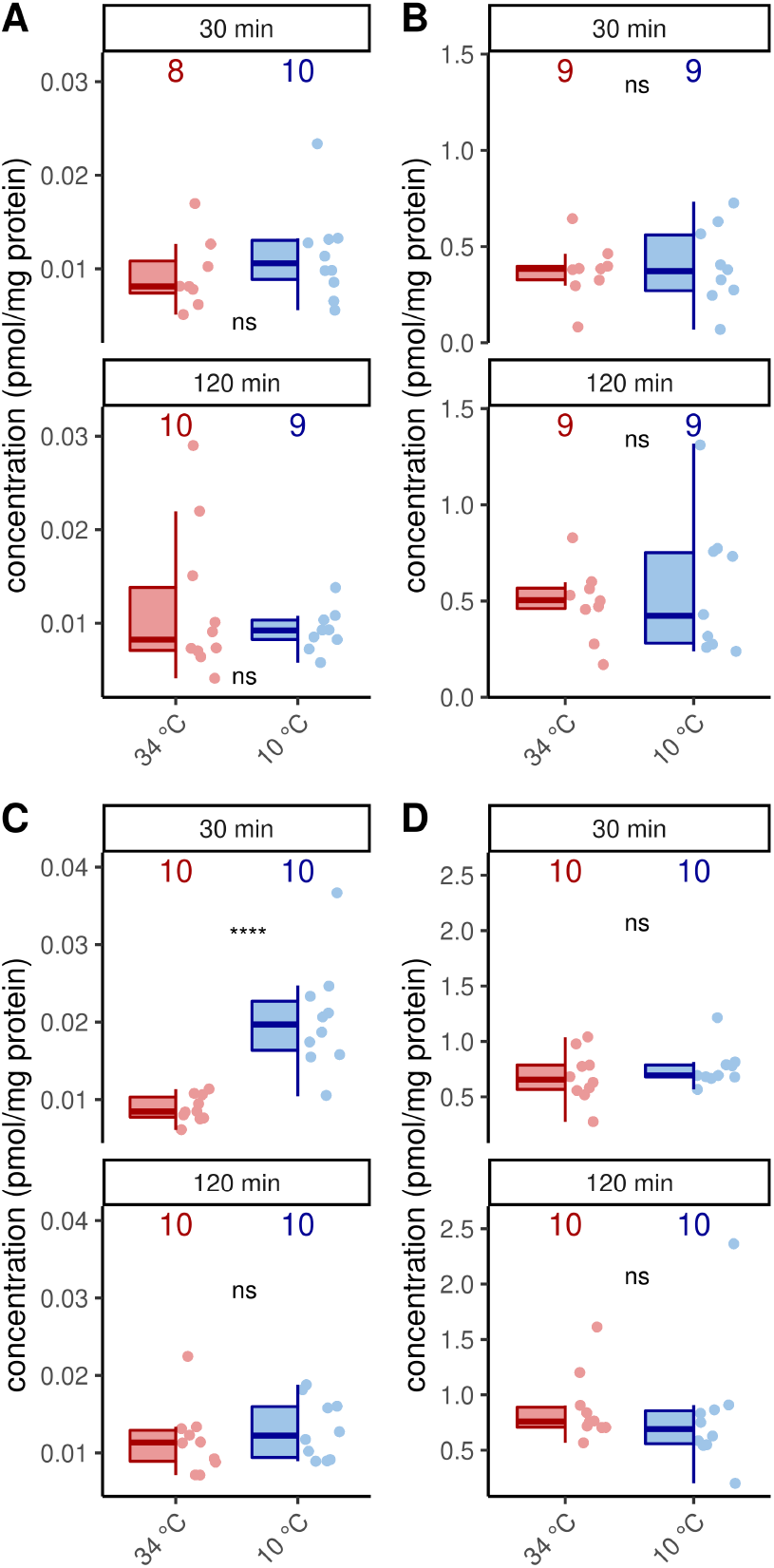
Octopamine concentration are stable under cold stress conditions. Only flight muscles after 30 min cold stress show increased octopamine concentrations (p = 0.00008, *U* = 8, *z* = -3.96, *r* = -0.89, Mann-Whitney *U* test). The remaining analysis reveal no differences in the flight muscles (A, C) nor the MMTG (B, D) in one week old workerbees (A-B) or forager bees (C-D). ns = p *≥* 0.05, Mann-Whitney *U* test. For each group/data-set median *±* IQR (left part) and individual data points (right part) are shown.

### Octopamine injections cannot simulate cold stress

Octopamine is provided to the flight muscles at constant levels in order to prevent chill coma. Very likely, this octopamine is released to the flight muscles due to its essential role in thermogenesis. We finally wanted to know whether an octopamine signal could cause effects similar to cold stress. Octopamine injections into flight muscles did not increase RNA_total_ levels regardless of incubation time (Figure 5A). Similarly, octopamine receptor gene expression is not changed after 30 min (Figure 5B). However, after 120 min expression of *AmOARα1* and *AmOARβ1* is increased and expression of *AmOARβ34* is decreased (Figure 5B). *AmOARβ2* expression remains unchanged (Figure 5B). The same is true for most isoform of this gene (Figure 5C). Here, the only exception is *AmOARβ2×3* whos abundance decreases 30 min after octopamine injection (Figure 5C).

**Figure 5.**
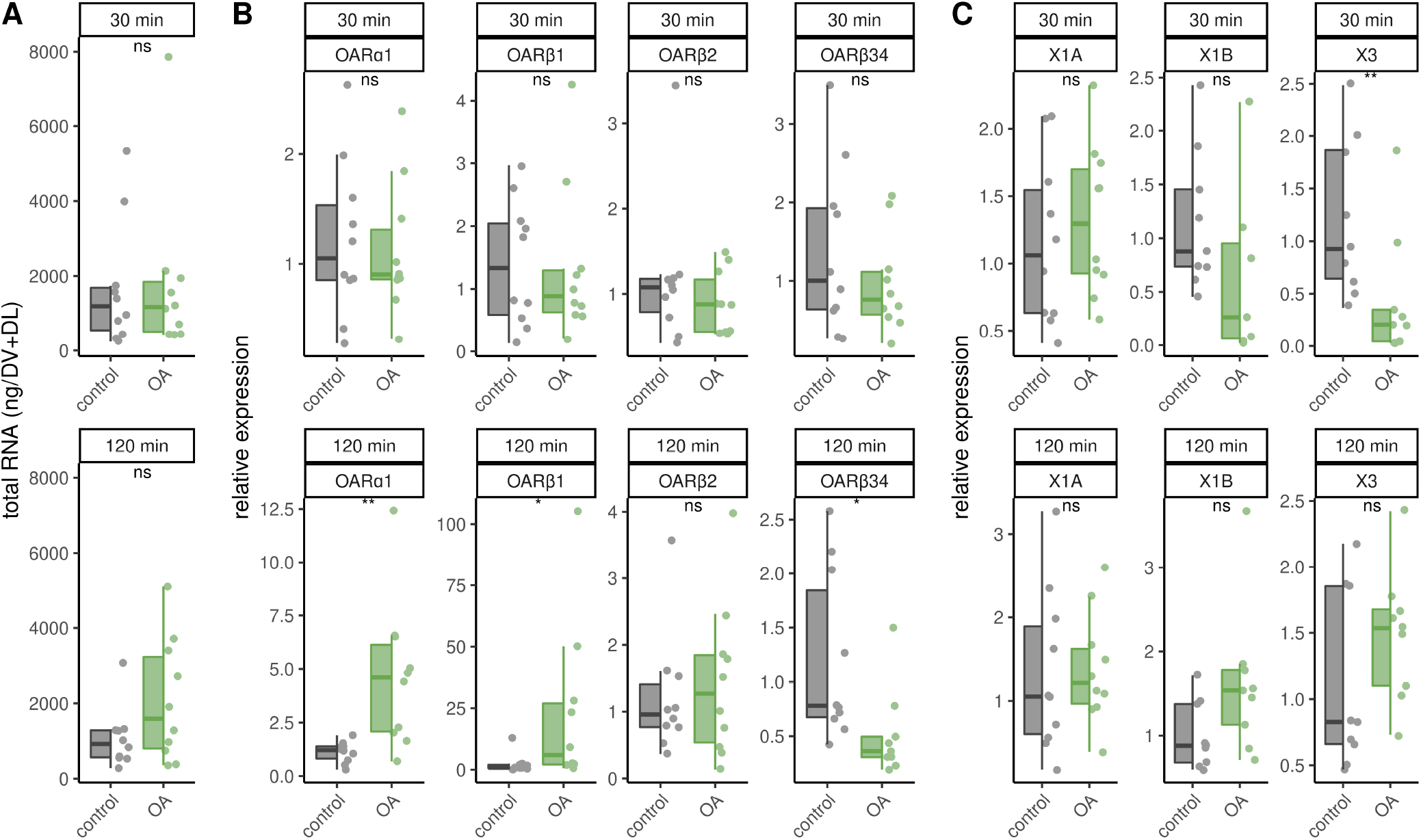
Effects of octopamine injections. Workerbees received octopamine injections in their flight muscles. After 30 and 120 min we measured RNA_total_ concentration **(A)**, octopamine receptor gene expression **(B)** and *AmOARβ2* isoform abundance **(C)**. * = p *<* 0.05, ** = p *<* 0.01, ns = p *≥* 0.05, Mann-Whitney *U* test. For each group/data-set*±* median IQR (left part) and individual data points (right part) are shown. For detailed statistics please see Table 3.

## DISCUSSION

Deviations from the optimal thermal state of a homoiothermic organism such as humans can have pathophysiological effects that, if they persist, lead to death (Obermeyer et al., 2017). Honeybees avoid comparable thermal stress induced consequences (Himmer, 1932; Weiss, 1962; Tautz et al., 2003; Groh et al., 2004; Wang et al., 2016) by maintaining thermostasis inside their colony as a super-organism (Simpson, 1961; Seeley and Visscher, 1985; Bujok et al., 2002; Stabentheiner et al., 2003; Buckley et al., 2015; Stabentheiner et al., 2021). We could simulate this under laboratory conditions with a small number of workerbees that manage to survive two hours of cold stress by effectively performing thermogenesis. This approach allowed us to analyze how the thoracic neuro-muscular system responds to cold stress at two distinct but essential levels (octopamine concentrations, expression of receptor genes) and to test the hypothesis whether this system is maintained at a constant level.

**Table 3.** Results of the statistical analysis (Mann-Whitney *U* test) of the effects of octopamine injections in to workerbee flight muscless. For visualization please see Figure 5.

| time | subject | $N_{control}$ | $N_{OA}$ | $U$ | $p$ | $z$ | $r$ | |
| --- | --- | --- | --- | --- | --- | --- | --- | --- |
| 30 min | $RNA_{total}$ | 10 | 10 | 48.00 | 0.91 | -0.11 | -0.02 | ns |
| 120 min | $RNA_{total}$ | 10 | 10 | 34.00 | 0.25 | -1.16 | -0.26 | ns |
| 30 min | *AmOARα1* | 10 | 10 | 53.00 | 0.85 | -0.19 | -0.04 | ns |
| 30 min | *AmOARβ1* | 10 | 10 | 53.00 | 0.85 | -0.19 | -0.04 | ns |
| 30 min | *AmOARβ2* | 10 | 10 | 56.00 | 0.68 | -0.41 | -0.09 | ns |
| 30 min | *AmOARβ34* | 10 | 10 | 58.00 | 0.58 | -0.55 | -0.12 | ns |
| 120 min | *AmOARα1* | 10 | 10 | 9.00 | 0.001 | -3.28 | -0.73 | ** |
| 120 min | *AmOARβ1* | 10 | 10 | 17.00 | 0.01 | -2.53 | -0.57 | * |
| 120 min | *AmOARβ2* | 10 | 10 | 46.00 | 0.80 | -0.26 | -0.06 | ns |
| 120 min | *AmOARβ34* | 10 | 9 | 76.00 | 0.01 | -2.57 | -0.59 | * |
| 30 min | *AmOARβ2X1A* | 10 | 10 | 40.00 | 0.48 | -0.70 | -0.16 | ns |
| 30 min | *AmOARβ2X1B* | 9 | 7 | 46.00 | 0.14 | -1.47 | -0.37 | ns |
| 30 min | *AmOARβ2X3* | 9 | 9 | 70.00 | 0.008 | -2.66 | -0.63 | ** |
| 120 min | *AmOARβ2X1A* | 10 | 10 | 43.00 | 0.63 | -0.48 | -0.11 | ns |
| 120 min | *AmOARβ2X1B* | 9 | 9 | 18.00 | 0.05 | -1.96 | -0.46 | ns |
| 120 min | *AmOARβ2X3* | 9 | 9 | 26.00 | 0.22 | -1.22 | -0.29 | ns |

First of all, this requires the supply of sufficient amount of octopamine. In fact, we found no cold stress associated differences in octopamine levels in the MMTG and in the flight muscles. With the exception of forager bees, where the octopamine level increased in both flight muscles after 30 min but returned to equilibrium after 120 min. The MMTG harbors flight muscle innervating octopaminergic neurons, but if octopamine is not present in sufficient quantities or if binding to its receptors is prevented, thermogenesis is compromised (Kaya-Zeeb et al., 2022). Unfortunately, with our method we cannot quantify octopamine in real-time and we cannot distinguish between vesicular and released octopamine. Consequently, we do not know how much octopamine is released per release event, how often such events occur, how long it remains at the target site and if released octopamine is recycled effectively. Especially the latter would probably be an important mechanism to deal efficiently with limited resources under extreme conditions (Schroeder and Jordan, 2012) and simultaneously maintain flight muscle functionality. Future studies could address these issues by combining different methods, which may include electrochemical microsensors (Phillips and Wightman, 2003; Jarriault et al., 2018), electrophysiological recordings (Ting and Phillips, 2007) and molecular and functional analyses of honeybee monoamine transporters (Torres et al., 2003; Zhang et al., 2019).

Similar to octopamine concentrations, the overall octopamine receptor gene expression does not appear to be strongly affected by cold stress. This is remarkable because a cold stress-induced increase in the amount of flight muscle RNA_total_ points to an increased transcription efficiency and reflects the increased need for the provision of specific newly synthesized proteins. The fact that octopamine receptor gene expression is not delayed in this process underscores the importance of octopamine signaling to the flight muscles and suggests that receptors have turnover rates that require some degree of re-synthesis of receptor proteins if their function is to be maintained. The majority of G-Protein coupled receptors (GPCRs) owns ligand-activated mechanisms that cause an arrestin-mediated removal of the receptor protein from the plasma membrane (Lefkowitz and Shenoy, 2005; Kelly et al., 2008; Tobin et al., 2008) which is also true for octopamine receptors (Hoff et al., 2011). Internalized GPCRs can either be subjected to membrane reintegration or degradation. If the latter is the case in the flight muscles, *de novo* protein synthesis is required to maintain thermogenesis functionality.

This seems to be important for *AmOARβ2*, which apparently is subject to complex alternative splicing. However, only *AmOARβ2×1A* seems to be associated with cold stress. This isoform encodes a functional described β octopamine receptor (Balfanz et al., 2013), which, according to our previous results, is the most likely octopamine receptor subtype in the service of thermogenesis (Kaya-Zeeb et al., 2022). The flight muscle abundance of *AmOARβ2×1A* is increased in one week old bees after 30 and 120 min of cold stress, even if the total expression of the *AmOARβ2* gene shows the same tendency only after two hours. The flight muscle expression of *AmOARβ2* increases with age (Kaya-Zeeb et al., 2022) and based on our new results, we suspect that a fast increase is possible if needed. Accordingly, forager bees which are usually older than three weeks (Winston, 1991) should have higher *AmOARβ2* expression levels. Here, cold stress does not elicit changes in *AmOARβ2* expression or *AmOARβ2×1A* abundance. The remaining isoforms do not follow a consistent trend over time (one week old bees) or are down-regulated after two hours (forager bees). For now, it is unlikely that these isoforms are important for thermogenesis, and future studies need to decipher their functions. This is also true for the remaining octopamine receptor genes, since we were not able to induce consistent expression changes by cold stress. We assume that they have important functions in flight muscle cells or associated tissues (e.g. nerves, trachea) as shown in flies (Sigrist and Andlauer, 2011; El-Kholy et al., 2015; Sujkowski et al., 2017, 2020). However, most likely they are not directly related to cold stress and thermogenesis.

Previously we could show that octopamine injections mimic cold stress induced *AmGAPDH* expression increase (Kaya-Zeeb et al., 2022). However, octopamine injections could not mimic cold-stress induced effects on RNA_total_ and *AmOARβ2* expression (including splicing). Furthermore, the changes in expression of the remaining octopamine receptor genes induced by the injections after two hours are probably nonspecific side effects, because comparable results are not observed under cold stress conditions. Thus, additional factors such as other neuromodulators (Salvador et al., 2021), hormones (Corona et al., 2007) or myokines (Schoenfeld, 2013; Pedersen et al., 2007; Pedersen, 2013) must be considered. It will be exciting to determine their properties and whether and how they coordinate with the octopaminergic system to maintain flight muscle function under cold stress.

Since the thoracic neuro-muscular octopaminergic system has an essential role in regulating thermogenesis, we expected a response to extreme temperature conditions. Ultimately, there are two plausible possibilities. The first option involves the up- and down-regulation of certain parts of the signaling cascade as required (Hadcock and Malbon, 1988a,b; Kim et al., 2003). Alternatively, critical components can be maintained on a constant synthesis and turnover rate (steady-state) to enable continuous operational readiness (Brodie et al., 1966; Kim et al., 2003). To date, our data suggest that the latter is the case. Constant octopamine concentrations indicate that octopamine can be released to the flight muscles if necessary, whether under optimal thermal conditions or under cold stress when thermogenesis is at full load. The same is true for octopamine receptor expression. This behavior of the thoracic neuro-muscular octopaminergic system makes perfect sense, since honeybees rely heavily on their abilities to regulate temperature (Stabentheiner et al., 2003, 2010). If they are not immediately available or even fail, the survival of the colony is very likely to be at risk. In the face of progressive climate change, the capability of thermostasis becomes increasingly important. The ability to maintain important physiological processes during increasingly frequent extreme weather events with sharply falling or rising temperatures represents a major challenge for all living organisms. Almost all relevant physiological processes are subject to a strong temperature dependence in their reaction rate (Hegarty, 1973; Reyes et al., 2008). Honeybees seem to be well equipped for such challenges and thermogenesis (and with this octopamine signaling) seems to play a crucial role here.

The question arises whether and how long the system can be kept functional if critical resources are scarce or no longer available. Our results show that one week old workerbees provided with only a carbohydrate source, but not amino acids or proteins, cannot maintain thermogenesis. Bees at this age are capable of thermogenesis (Stabentheiner et al., 2010). However, in our case they will lack essential amino acids (de Groot, 1953), which are required for the synthesis of octopamine and other monoamines (Roeder, 1999; Blenau and Thamm, 2011) as well as proteins (Cremonz et al., 1998). In addition, a possible deficiency of essential fatty acids, vitamins and electrolytes must be considered (Zarchin et al., 2017; Gribakin et al., 1987). Although these bees also respond to cold stress with a massive up-regulation of transcription in their flight muscles, they are incapable of effective thermogenesis and fall into a chill coma. It is conceivable that there is not enough octopamine available for thermogenesis or that the flight muscle is not optimally developed, or a combination of both. Future experiments need to show how certain aspects of nutrition affect the steady-state of the thoracic neuro-muscular octopaminergic system and how limited resource availability may disrupts thermoregulation at both the individual and colony level. This could become apparent if natural food resources are increasingly decimated by the ongoing intensification of agricultural cultivation) (Rich and Woodruff, 1996; Biesmeijer et al., 2006; Vaudo et al., 2015).

With this study, we provide an important contribution to the better understanding of the fundamental and essential physiological response of worker honeybees to cold stress. Moreover, we provide an important basis for comparative analyses with species that are not as resistant to temperature stress as honeybees. To capture both extreme possibilities of extreme weather events, future studies should definitely include heat stress. Continuing these analyses will help us gain a more comprehensive picture of strategies for adapting to changing environmental conditions, whether in the context of natural evolutionary processes or in the course of global change.

## ACKNOWLEDGMENTS

We thank Dirk Ahrens-Lagast for bee keeping; Johannes Spaethe, Ricarda Scheiner and Petra Högger for supplying technical equipment. A special thanks goes to Christian Wegener, Flavio Roces, Johannes Spaethe, Felix Schilcher and Ricarda Scheiner for fruitful discussions and helpful comments.

This work was supported by a grand of the Deutsche Forschungsgemeinschaft to MT (TH2264/2-1).

## SUPPLEMENTAL DATA

**Table S1.** Results of the statistical analysis (Mann-Whitney *U* test) of dopamine (DA) and serotonin (5-HT) quantification in workerbee flight muscles (DV+DL) and mesometathoracic ganglion (MMTG) under cold stress. Serotonin was only detectable in the MMTG but not the flight muscles. Tyramine was not quantifiable in many samples and was therefore not considered. For visualization see Figure S1.

| bees | tissue | time | amine | $N_{34^{\circ}\text{C}}$ | $N_{10^{\circ}\text{C}}$ | $U$ | $p$ | $z$ | $r$ |
| --- | --- | --- | --- | --- | --- | --- | --- | --- | --- |
| one week old | DV+DL | 30 min | DA | 8 | 10 | 26.00 | 0.24 | -1.18 | -0.28 |
| one week old | DV+DL | 120 min | DA | 10 | 9 | 45.00 | 1.00 | 0.00 | 0.00 |
| one week old | MMTG | 30 min | DA | 9 | 9 | 50.00 | 0.44 | -0.78 | -0.18 |
| one week old | MMTG | 30 min | 5-HT | 9 | 9 | 55.00 | 0.22 | -1.22 | -0.29 |
| one week old | MMTG | 120 min | DA | 9 | 9 | 38.00 | 0.86 | -0.17 | -0.04 |
| one week old | MMTG | 120 min | 5-HT | 7 | 9 | 47.00 | 0.11 | -1.58 | -0.40 |
| forager | DV+DL | 30 min | DA | 10 | 10 | 37.00 | 0.35 | -0.93 | -0.21 |
| forager | DV+DL | 120 min | DA | 10 | 10 | 48.00 | 0.91 | -0.11 | -0.02 |
| forager | MMTG | 30 min | DA | 10 | 10 | 48.00 | 0.91 | -0.11 | -0.02 |
| forager | MMTG | 30 min | 5-HT | 10 | 10 | 36.00 | 0.32 | -1.00 | -0.22 |
| forager | MMTG | 120 min | DA | 10 | 10 | 76.00 | 0.05 | -1.94 | -0.43 |
| forager | MMTG | 120 min | 5-HT | 8 | 6 | 31.00 | 0.41 | -0.82 | -0.22 |

**Figure S1.**
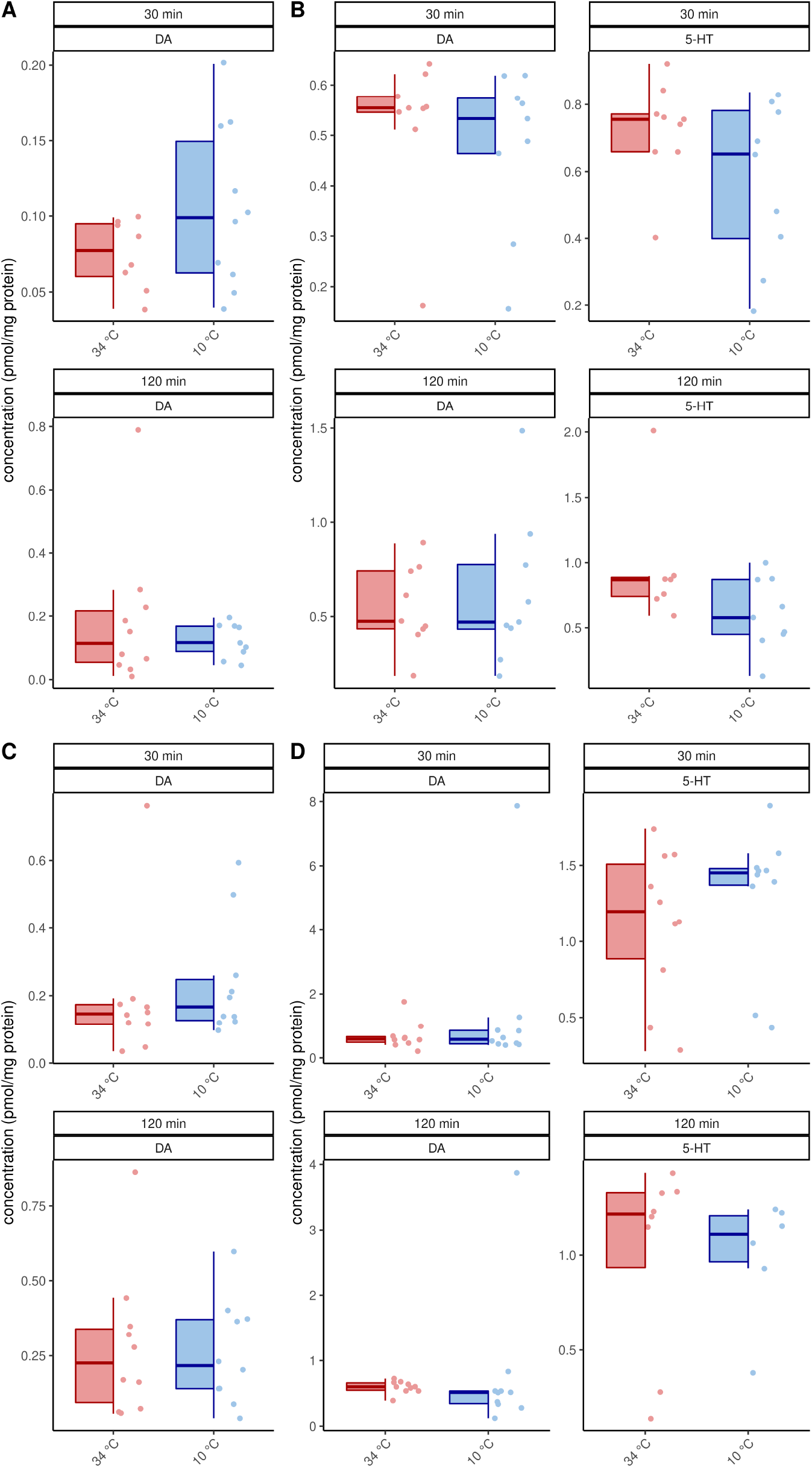
Dopamine (DA) and Serotonin (5-HT) concentrations in the flight muscles (A, C) and the MMTG (B, D) under cold stress. For statistics see Table S1. For each group/data-set median *±* IQR (left part) and individual data points (right part) are shown.

